# MicroRNA signatures of equine asthma endotypes in serum and bronchoalveolar lavage fluid

**DOI:** 10.64898/2026.03.29.715111

**Authors:** Emilie Rudolph Røgild, Emilio Mármol-Sánchez, Katrine Toft, Sanni Hansen, Susanna Cirera

**Affiliations:** Department of Veterinary and Animal Sciences, University of Copenhagen, Copenhagen, Denmark; Center for Evolutionary Hologenomics, The Globe Institute, University of Copenhagen, Copenhagen, Denmark; Department of Veterinary Clinical Sciences, University of Copenhagen, Copenhagen, Denmark

## Abstract

Equine asthma (EA) is a highly prevalent, chronic, inflammatory disease of the lower airways ranging from mild-to-moderate to severe clinical presentations. Diagnosis currently relies on bronchoalveolar lavage fluid (BALF) cytology, an invasive method associated with interobserver variability, which highlights the need for more reproducible approaches. MicroRNAs (miRNAs) are small noncoding RNAs involved in post-transcriptional gene regulation. They are stable and readily detectable in body fluids and have shown promising results as biomarkers in human asthma. The aim of this study was to characterize miRNA abundance profiles in BALF and serum from horses with distinct EA endotypes to evaluate their biomarker potential and explore their involvement in disease pathogenesis. A total of 43 horses were included and classified as either EA (n=32) or controls (n=11), based on clinical examination and BALF cytology. The EA horses were further divided into three endotypes based on BALF inflammatory cell composition: neutrophilic asthma (n=10), mastocytic asthma (n=15), and mixed asthma (n=7). RNA was isolated from both serum and BALF samples and analyzed by quantitative real-time PCR (qPCR) targeting 103 miRNAs linked to asthma and pulmonary inflammation in humans. Differential miRNA abundance was analyzed across EA endotypes. The most significantly differentially abundant miRNAs were used for *in silico* target prediction and pathway enrichment analyses. Horses with mixed EA had significantly lower levels of eca-miR-125a-3p and eca-miR-125b-5p in BALF compared to controls. Additionally, eca-miR-146a-5p abundance was significantly increased in BALF from horses with neutrophilic EA compared to mastocytic EA. Target and pathway enrichment analyses for eca-miR-146a-5p identified immune-relevant pathways, such as MAPK and T-cell receptor signaling, supporting its involvement in inflammatory processes associated with asthma. This study identified three promising candidates, eca-miR-125a-3p, eca-miR-125b-5p, and eca-miR-146a-5p, as potential biomarkers associated with different EA endotypes. These miRNAs are interesting candidates for further investigation in an independent cohort.

## Introduction

Equine asthma (EA) is a naturally occurring, chronic, non-infectious inflammatory disease of the lower airways [1]. EA is broadly categorized into severe (sEA) and mild-to-moderate (mEA) phenotypes [2]. Severe cases typically affect older horses that present with marked respiratory effort at rest, whereas mild-to-moderate cases manifest primarily as chronic coughing and reduced performance. The mEA phenotype can further be subdivided into endotypes based on the percentage of neutrophils, mast cells and eosinophils or a combination of these in the bronchoalveolar lavage fluid (BALF) [3]. Clinically and pathophysiologically, EA shares important features with human asthma, including airway inflammation, mucus hypersecretion, bronchial hyperresponsiveness, and variable airflow obstruction [4]. Despite years of research, the pathogenesis of EA remains incompletely understood, and current diagnostic and therapeutic strategies are consequently suboptimal. The EA diagnosis relies on clinical examination combined with bronchoalveolar lavage fluid (BALF) cytology, which is considered the gold standard for assessing lower airway inflammation [1,2,5,6]. However, BALF classification based solely on cytology may not fully capture underlying molecular endotypes [1,7,8]. Consequently, there is a need for investigations into complementary biomarkers that can enhance understanding of EA pathophysiology, improve phenotyping and possibly provide less invasive diagnostic options [1,9].

MicroRNAs (miRNAs) are small, stable noncoding RNAs that regulate gene expression post-transcriptionally and play critical roles in immune modulation, inflammation, and tissue remodeling [10]. These molecules are present in biofluids, including serum and BALF, often encapsulated in extracellular vesicles, and have been implicated in key pathogenic pathways in human asthma, such as Th2 differentiation, eosinophilic inflammation, epithelial dysfunction, and airway remodeling [11]. Several studies have identified miRNA signatures associated with disease severity, inflammatory phenotype, and treatment response [11,12]. However, variability between studies and limited validation across independent cohorts have left uncertainties regarding their clinical utility. While no studies have profiled miRNAs in BALF of horses, previous reports on serum and plasma [13,14], bronchial smooth muscle [15], and lung tissue [12] suggest that miRNAs could provide novel insights into equine EA disease mechanisms and support more precise phenotyping. Given the shared pathophysiology between equine and human asthma, the investigation of miRNAs in EA could not only improve diagnostics in horses but also provide translational insights into human airway disease. The aim of the present study was to characterize miRNA profiles in serum and BALF from horses with different EA BALF cytology profiles, in order to evaluate their biomarker potential for disease discrimination and improve understanding of molecular mechanisms in EA pathogenesis.

## Materials and methods

### Study population

A total of 43 horses were enrolled in this cross-sectional observational study, with serum and BALF samples collected from each horse upon admission to the Teaching Hospital for Large Animals in Taastrup between 30 October 2023 and 20 March 2025 (see **Supplementary Table 1** for horse characteristics). This study was approved by the ethics committee of the Department of Veterinary Clinical Sciences, University of Copenhagen (2020-013) with informed client consent.

Diagnosis of EA was established based on a combination of owner-reported anamnesis (cough or poor performance), clinical signs (respiratory rate and type, lung auscultation and/or spontaneous cough during the clinical examination) and BALF cytology. Reference intervals for the EA classification applied to BALF cytology samples were: >10% neutrophils, >5% eosinophils and/or >5% mast cells [2]. Classification of the horses into EA endotypes was based on the BALF cytology. The groups were defined as follows: neutrophilic EA (>10% neutrophils, <5% eosinophils and <5% mast cells), mastocytic EA (<10% neutrophils, <5% eosinophils and >5% mast cells), mixed EA (>10% neutrophils, <5% eosinophils and >5% mast cells), and controls (<10% neutrophils, <5% eosinophils and <5% mast cells) [2]. Horses were included as controls if they were admitted for a full diagnostic workup for poor performance, had no history of coughing, showed BALF cytology within the normal reference range, and received a final diagnosis not related to lower airway disease.

### Endoscopic examination and bronchoalveolar lavage fluid extraction

A medical history was obtained from the owner of each horse. A clinical examination, including evaluation of the horse under saddle or during lunging, was performed prior to the endoscopic examination. Blood samples were collected in serum separator tubes (BD, Franklin Lakes, NJ, USA) using a vacutainer system. Subsequently, all horses were sedated with a combination of detomidine (Domosedan 10 mg/mL, Orion Pharma, Denmark) 0.01 mg/kg IV and butorphanol (Dolorex, MSD Animal Health) 0.01 mg/kg IV and positioned in a stock. A 3000 mm, 10.4 mm diameter endoscope (Karl Storz 60 130 PKS G28-300, Germany) was inserted via the ventral meatus through the nasopharynx and trachea. The endoscope was advanced into the right stem bronchus until wedged in a caudo-dorsal bronchus. A total of 250 mL sterile isotonic saline (0.9%, Braun, Germany) was infused and immediately aspirated. Eight mL of the aspirate was transferred to EDTA tubes for cytology, and the remaining BALF was transferred to 150 mL sterile bottles for miRNA processing.

### Bronchoalveolar lavage fluid cytology

For all samples, 200 μL of EDTA-stabilized BALF was applied to the funnel of a cytospin and centrifuged at 93 × g for 8 minutes (StatSpin Cytofuge 12, USA). The cytospin smears were then fixed in ethanol and stained with May-Grünwald-Giemsa (Merck, Germany) prior to the microscopic examination, using a 40× magnification microscope (Leica Microsystems, dm1000). To obtain a BALF differential cell count, the proportion of macrophages, lymphocytes, neutrophils, mast cells, and eosinophils was calculated for each cytospin by counting a total of 200 cells.

### Preprocessing of serum and bronchoalveolar lavage fluid samples for miRNA analysis

Blood samples were centrifuged at 800 × g for 10 minutes within 1-2 hours after collection and the serum fraction thereafter stored at −70°C until RNA isolation. The BALF samples were first concentrated from 15 mL to 500 µL as follows: Initially, BALF samples were centrifuged at 3000 × g for 10 minutes at 4°C and the supernatant of each sample was filtered through a 0.2 µm filter (Millipore) into a 50 mL tube. The filtrate was then stored at −70°C until further purification. Following thawing on ice, 15 mL of total BALF filtrate was re-filtered through a 0.2 µm filter and processed through a 100 kDa filter (Amicon) by centrifuging at 300 × g at 4°C until the residual volume in the filter was reduced to approximately 500 µL. The resulting concentrated fluid was then transferred to a 2 mL cryotube and stored at −70°C until RNA isolation.

### RNA isolation and cDNA synthesis

Total RNA was isolated from both serum and BALF samples using the miRNeasy serum/plasma kit (Qiagen) following the manufacturer’s protocol. Briefly, 200 µL of serum and 500 µL of BALF were mixed with QIAzol reagent, followed by phase separation using chloroform and RNA purification on RNeasy MinElute spin columns. The isolated RNA molecules were then eluted in RNase-free water and quantified using a NanoDrop spectrophotometer (ThermoFisher Scientific). All RNA samples were then stored at −80°C until further analysis. Fifty ng of RNA from each BALF and serum sample was reverse transcribed into cDNA [16,17]. Two replicates for each RNA sample were then synthesized, as well as two negative controls without including Poly(A) polymerase (-PAP) in the reaction for each sample type (BALF and serum).

### MicroRNA selection and primer design

The miRNAs to be profiled were selected based on literature search using “microRNAs in asthma” as key words. The list of miRNAs initially included in this study (n=103), and their mature sequences acquired from miRBase database [18] are available in **Supplementary Table 2**. Each mature miRNA sequence was used for primer design using the miRprimer software [19]. A list of forward and reverse primers for each miRNA used in this study is also available in **Supplementary Table 2**.

### Quantitative real-time PCR

All miRNA assays were first evaluated and qualified using a QuantiStudio3 qPCR system (Applied Biosystems by ThermoFisher) and the PowerUp Master Mix (ThermoFisher) according to manufacturer’s instructions. The list of assays that qualified and were therefore further processed (n=49) is available in **Supplementary Table 3.** Subsequently, the optimal number of preamplification cycles and the number of points in the standard curve were determined using a FlexSix IFC chip (Fluidigm) according to manufacturer’s protocol. Following optimization, the complete qPCR experiment was performed using two 96×96 IFC chips on the BioMark™ HD system (Fluidigm) — one chip for BALF samples, and one for serum samples, respectively. All cDNA samples were pre-amplified for 20 cycles, exonuclease treated and diluted 8× in TE buffer prior to running in two 96×96 IFC chips in the BioMark™ HD system following standard Fluidigm protocols. The corresponding standard curve, the two -PAP controls and a reaction without cDNA (NTC) were also included in each chip.

### Data curation and preprocessing

Following qPCR, all assays and samples were visually examined using the Fluidigm software. Samples showing atypical amplification plots or melting curves were excluded. Assays with amplification in NTC and/or -PAP controls in close range of those from experimental samples, as well as with missing quantification cycle values (Cq) were also excluded. PCR efficiency was calculated from the slope of the standard curve and assays in the range of 80-110% PCR efficiency were accepted. Lastly, sample replicates were excluded if the Cq value differed substantially from the other replicate within the same assay. Subsequently, manually curated data was imported into GenEx^TM^ software v6.0 (Multid) for processing. BALF and serum samples were processed separately. Briefly, samples were first corrected for PCR efficiency. Reference miRNAs were then determined using the NormFinder algorithm [20]. For BALF samples, eca-let-7d-5p, eca-miR-27a-3p, and eca-let-7a-5p were selected as normalizers, while eca-miR-27b-3p, eca-miR-30c-5p, and eca-miR-21-5p were chosen for serum samples. Using the three selected stably abundant assays for BALF and serum, efficiency-corrected raw qPCR values were then normalized. Subsequently, sample replicates were averaged, and relative quantities were calculated scaling to the sample with the lowest abundance per each assay. All data were finally log_2_-transformed prior to statistical analysis (see **Supplementary Table 4** for serum and **Supplementary Table 5** for BALF).

### Statistical analysis

Significant differences in the average abundance of each miRNA profiled across control horses, and horses with mastocytic EA, mixed EA or neutrophilic EA were assessed using the limma empirical Bayes procedure from limma v3.60.4 R package [21]. Sex, age, and castration status (**Supplementary Table 1**) were introduced as covariates when fitting the linear mixed model prior to the use of the empirical *ebayes* function. Missing values for age were imputed with the average of all other horses in the study. Breed was excluded as a covariate due to the high number of levels relative to sample size (17 breeds across 43 sampled horses), with many breeds only represented by as few as 1 individual. Missing values for qPCR miRNA assays were imputed by calculating the average abundance across all samples for a given assay. The obtained *P*-value estimates were corrected for multiple testing using the False Discovery Rate (FDR) method [22]. The miRNA assays that passed pre-processing (36 for serum, and 35 for BALF) included for differential abundance analyses are listed in **Supplementary Table 6**.

The following pairwise contrasts were considered: i) control vs. all types of EA combined, ii) control vs. mastocytic EA, iii) control vs. mixed EA, iv) control vs. neutrophilic EA, v) mastocytic EA vs. mixed EA, vi) mastocytic EA vs. neutrophilic EA, and vii) mixed EA vs. neutrophilic EA for serum and BALF, respectively. The threshold for significance was set at *P*-value < 0.05 and absolute FC > 1.5. MiRNA assays with FDR < 0.05 and absolute FC > 1.5 (n=3) were considered as strongly differentially abundant and used for further target prediction analyses. When control horses were used in a contrast, these were set as the baseline, meaning that any overabundance in EA samples would appear with a positive FC, or vice versa. For the contrast comparing mastocytic EA vs. mixed EA, mastocytic EA vs. neutrophilic EA, and mixed EA vs. neutrophilic EA, the former was set as the baseline. Fold changes and the average abundance for each miRNA assay in both groups for each contrast are shown as the base 2 antilog of the log_2_ (FC) values returned by limma, and as the log_2_-transformed abundance estimates used as input for statistical inference, respectively (see **Supplementary Table 7** for serum samples and **Supplementary Table 8** for BALF samples).

To test the association between significantly differentially abundant miRNAs across groups and cell composition measured in BALF cytospins, we calculated Pearson’s correlation coefficients (*r*). For this analysis we used the log_2_-normalized relative quantity values of selected miRNAs profiled in BALF and serum, and the quantification of cell composition, expressed in percentage values, obtained for BALF cytospins in each individual horse analyzed. A linear regression trend was fitted to each pairwise relationship with a 95% confidence interval.

### Target prediction and enrichment analyses

Putative mRNA gene targets of the miRNAs that showed strong significant differences (FDR < 0.05 and |FC| > 1.5) in any of the aforementioned contrasts after multiple testing correction were predicted using the SeedVicious v1.1 tool [23]. SeedVicious uses a simple nucleotide match among the miRNA seeds and 3’UTR sequences of target mRNAs to identify canonical miRNA-mRNA binding sites. We focused only on the 8mer matches given these types of miRNA-mRNA interactions are considered to be the strongest among the different types of known canonical ones [24]. The mature miRNA sequences of eca-miR-146a-5p, eca-miR-125a-3p, and eca-miR-125b-5p were obtained from miRBase database [18], while the 3’UTR sequences of all protein-coding mRNA genes annotated in the horse assembly (EquCab3.0) were obtained from Ensembl v115 [25] using the BioMart tool [26]. Only 3’UTR sequences longer or equal to 40 nucleotides were kept for target prediction analyses (see **Supplementary Table 9**). To assess shared biological functions among the predicted 8mer targets, the putative mRNA targets of eca-miR-146a-5p, eca-miR-125a-3p, and eca-miR-125b-5p (SeedVicious output) were analyzed using ShinyGO v0.85 [27]. This tool performs GO and pathway enrichment using a hypergeometric test with FDR correction [22]. The horse assembly (EquCab3.0) was selected as the reference species, and all annotated protein-coding mRNA genes used for prediction were defined as the background gene set. Default parameters were applied except for the minimum pathway size, which was set at 5. The KEGG database was used to identify enriched pathways. Ontologies were considered significant at FDR value < 0.05 (see **Supplementary Table 10**).

## Results

### Study population and clinical signs

The cohort was comprised of 28 geldings, 3 stallions, and 12 mares, representing a range of different breeds, with Icelandic horses being the largest single group (n = 5). Horses ranged in age from 2 to 20 years. Detailed information on age, sex, castration status, and breed for each horse is provided in **Supplementary Table 1**. Results from the owner-reported anamnesis as well as selected variables from the clinical examination are presented in **Table 1**.

**Table 1:** Owner-reported anamnesis, including coughing and/or poor performance, as well as body weight and body condition score (BCS) [28], are included in the table. Clinical examination findings, including rectal temperature, heart rate, respiratory rate and respiratory pattern, and the presence of coughing observed during the exercise component of the examination, are also provided.

|  | Control<br>(n=11) | Neutrophilic<br>EA (n=10) | Mastocytic<br>EA (n=15) | Mixed<br>EA (n=7) | Total<br>(n=43) |
| --- | --- | --- | --- | --- | --- |
| Owner reported |  |  |  |  |  |
| Cough (yes/total) | 0/11 | 8/10 | 12/15 | 5/7 | 25/43 |
| Poor performance (yes/total) | 0/11 | 7/10 | 13/15 | 7/7 | 27/43 |
| Clinical exam |  |  |  |  |  |
| Weight (kg) (mean ± SD) | 522 ± 99 | 581 ± 90 | 518 ± 116 | 448 ± 83 | 522 ± 106 |
| BCS (1-9) (mean ± SD) | 6.3 ± 1.2 | 6.1 ± 0.9 | 6.9 ± 1.4 | 5.3 ± 0.8 | 6.3 ± 1.2 |
| Rectal temperature (°C) (mean ± SD) | 37.7 ± 0.4 | 37.7 ± 0.3 | 37.9 ± 0.3 | 37.8 ± 0.3 | 37.8 ± 0.4 |
| Heart rate (beats/min) (mean ± SD) | 39 ± 8 | 41 ± 6 | 39 ± 6 | 39 ± 4 | 40 ± 6 |
| Respiratory rate (breaths/min) (mean ± SD) | 14 ± 3 | 22 ± 4 | 20 ± 8 | 20 ± 3 | 19 ± 6 |
| Respiratory type<br>(Thoracoabdominal/ abdominal) | 11/0 | 3/7 | 11/4 | 7/0 | 32/11 |
| Spontaneous cough during work<br>(yes/total) | 0/11 | 5/10 | 0/15 | 3/7 | 8/43 |

Abbreviations: EA: equine asthma; Control: control horses without equine asthma. SD = standard deviation. BCS: Body condition score [28].

### Bronchoalveolar lavage fluid cytology

Endotype classification was determined solely by inflammatory cell composition in BALF, resulting in three endotypes: neutrophilic EA (n = 10), mastocytic EA (n = 15), and mixed EA (n = 7), along with controls (n = 11). All horses within the mixed EA group had a combination of neutrophils and mast cells above reference intervals (**Figure 1**). See **Supplementary Table 1** for a comprehensive summary of percentages of each cell type in all BALF samples analyzed in this study.

**Figure 1:**
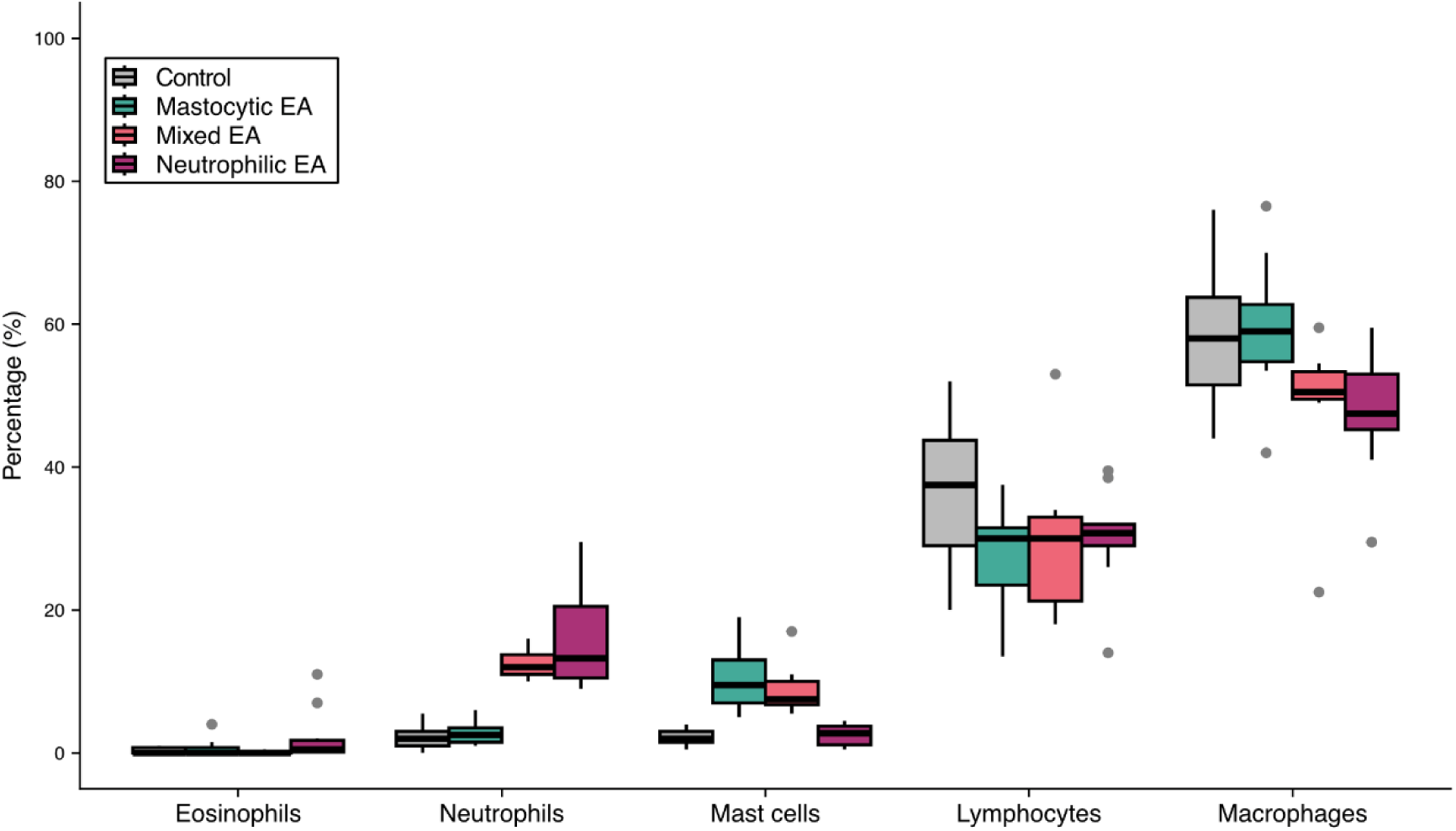
Distribution of the bronchoalveolar lavage fluid differential cell counts (%) including eosinophils, neutrophils, mast cells, lymphocytes and macrophages of the included horses (n=43), according to their equine asthma (EA) diagnosis (i.e. controls, and horses with mastocytic EA, mixed EA or neutrophilic EA). Percentage values for each analyzed horse and cell types are available in **Supplementary Table 1**.

### Differentially abundant microRNAs in serum

In serum samples, none of the analyzed miRNAs (**Supplementary Table 7**) were significantly differentially abundant between groups after FDR correction (FDR < 0.05, see **Table 2**). However, several miRNAs showed significantly differential abundance at nominal *P*-value < 0.05 and absolute fold change (FC) > 1.5.

**Table 2:** Differentially abundant microRNAs (miRNAs) in serum samples from control horses and horses with different equine asthma (EA) endotypes: *P*-value < 0.05; |Fold Change| > 1.5.

| Comparison | miRNA | Fold Change | $P$-value | FDR |
| --- | --- | --- | --- | --- |
| Control vs. all EA | eca-miR-200b-3p | 2.219 | 0.032 | 0.432 |
| Control vs. mastocytic EA | eca-miR-200b-3p | 2.278 | 0.049 | 0.889 |
| Control vs. mixed EA | eca-let-7a-5p | 1.588 | 0.015 | 0.386 |
|  | eca-miR-30d-5p | -1.757 | 0.029 | 0.386 |
|  | eca-miR-200b-3p | 2.796 | 0.043 | 0.386 |
| Control vs. neutrophilic EA | eca-miR-146a-5p | -2.008 | 0.026 | 0.488 |
|  | eca-miR-107b-3p | -1.809 | 0.036 | 0.488 |
| Mastocytic EA vs. neutrophilic EA | eca-miR-107b-3p | -2.006 | 0.01 | 0.315 |
|  | eca-miR-140-3p | -1.618 | 0.031 | 0.315 |
|  | eca-miR-145-5p | -1.636 | 0.034 | 0.315 |
| Mastocytic EA vs. mixed EA | - | - | - | - |
| Mixed EA vs. neutrophilic EA | eca-miR-107b-3p | -2.729 | 0.003 | 0.091 |
|  | eca-miR-27a-3p | -1.793 | 0.04 | 0.392 |

EA: equine asthma; FDR: false discovery rate.

Eca-miR-200b-3p showed consistently higher levels in serum from horses diagnosed with EA (considering all EA endotypes together) compared to controls, as well as when either mastocytic or mixed EA were compared to controls. The miRNA eca-let-7a-5p also showed increased serum abundance in horses with mixed EA compared to controls. On the contrary, eca-miR-30d-5p, eca-miR-107b-3p, and eca-miR-146a-5p showed lower levels in serum from horses with mixed EA and neutrophilic EA compared to controls, respectively. Furthermore, eca-miR-107b-3p, eca-miR-140-3p, eca-miR-145-5p, and eca-miR-27a-3p were significantly less abundant in horses with neutrophilic EA compared to either mastocytic or mixed EA respectively (see **Table 2**). A comprehensive report for differential abundance analyses in serum miRNAs profiled by qPCR is available in **Supplementary Table 7**.

To gain further insight into the abundance of the aforementioned miRNAs across all groups, a bar-plot was created (**Figure 2**), which illustrates the relative quantities of the significantly differentially abundant miRNAs in each group for the serum samples.

**Figure 2:**
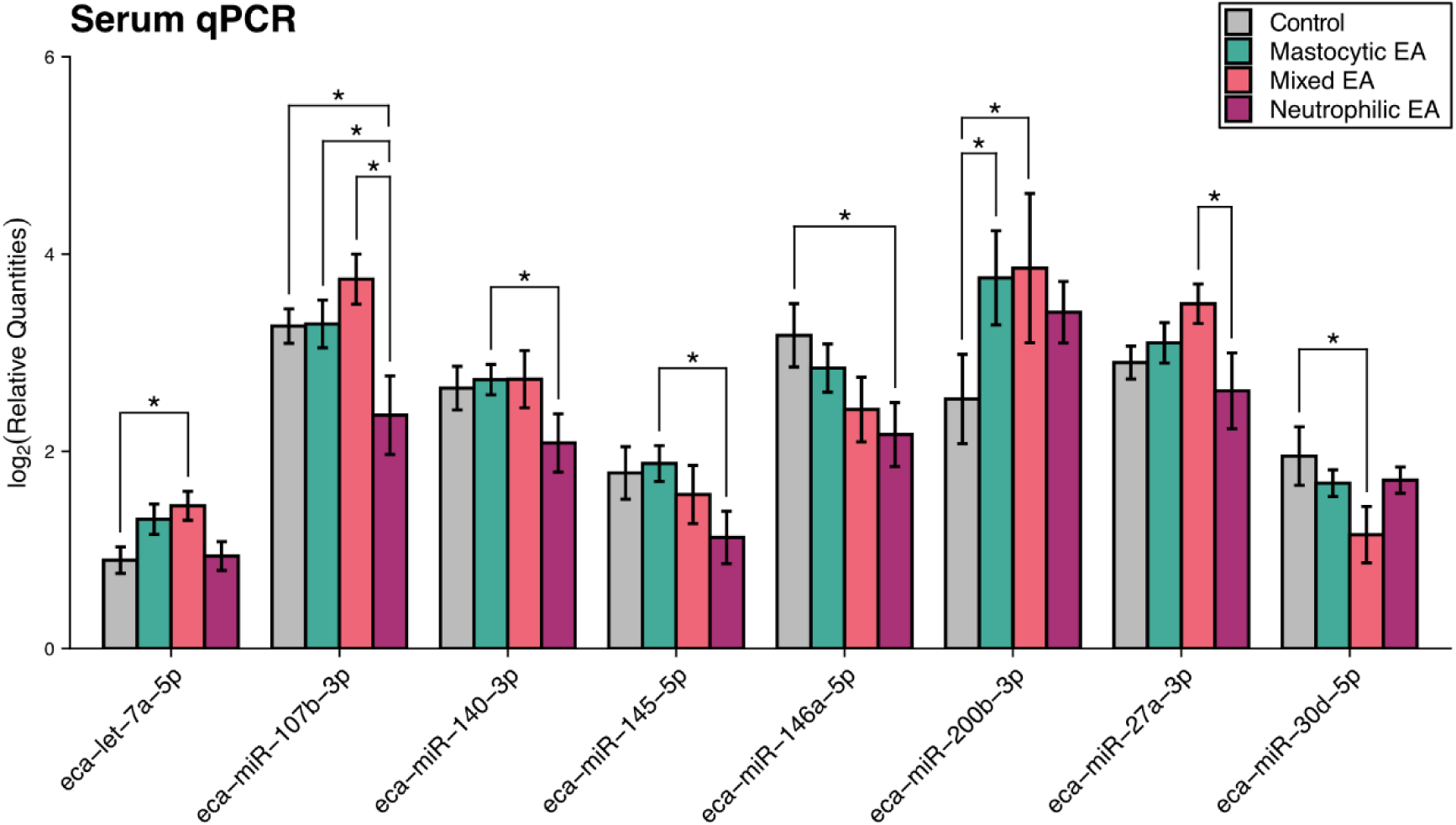
Relative quantities of significantly abundant miRNAs in serum samples from control horses and horses with different equine asthma (EA) endotypes. Significant differences at nominal level (|Fold Change| > 1.5; *P*-value < 0.05) are marked with a star (*).

### Differentially abundant microRNAs in bronchoalveolar lavage fluid

Analysis of the BALF samples revealed three miRNAs that were significantly differentially abundant between the groups after FDR correction (|FC| > 1.5; FDR < 0.05, see **Figure 3** and **Table 3** below), as well as other miRNAs that were significant at nominal level (|FC| > 1.5; *P*-value < 0.05). Two of the miRNAs that were highly significant after multiple testing correction were eca-miR-125a-3p and eca-miR-125b-5p, which were strongly downregulated in mixed EA horses compared to controls (**Table 3**). They were also significantly less abundant, albeit only at nominal level, in mixed EA horses compared to the neutrophilic EA group (see **Figure 3 and Table 3**). A full list for differential abundance analyses in BALF miRNAs is available in **Supplementary Table 8.**

**Figure 3:**
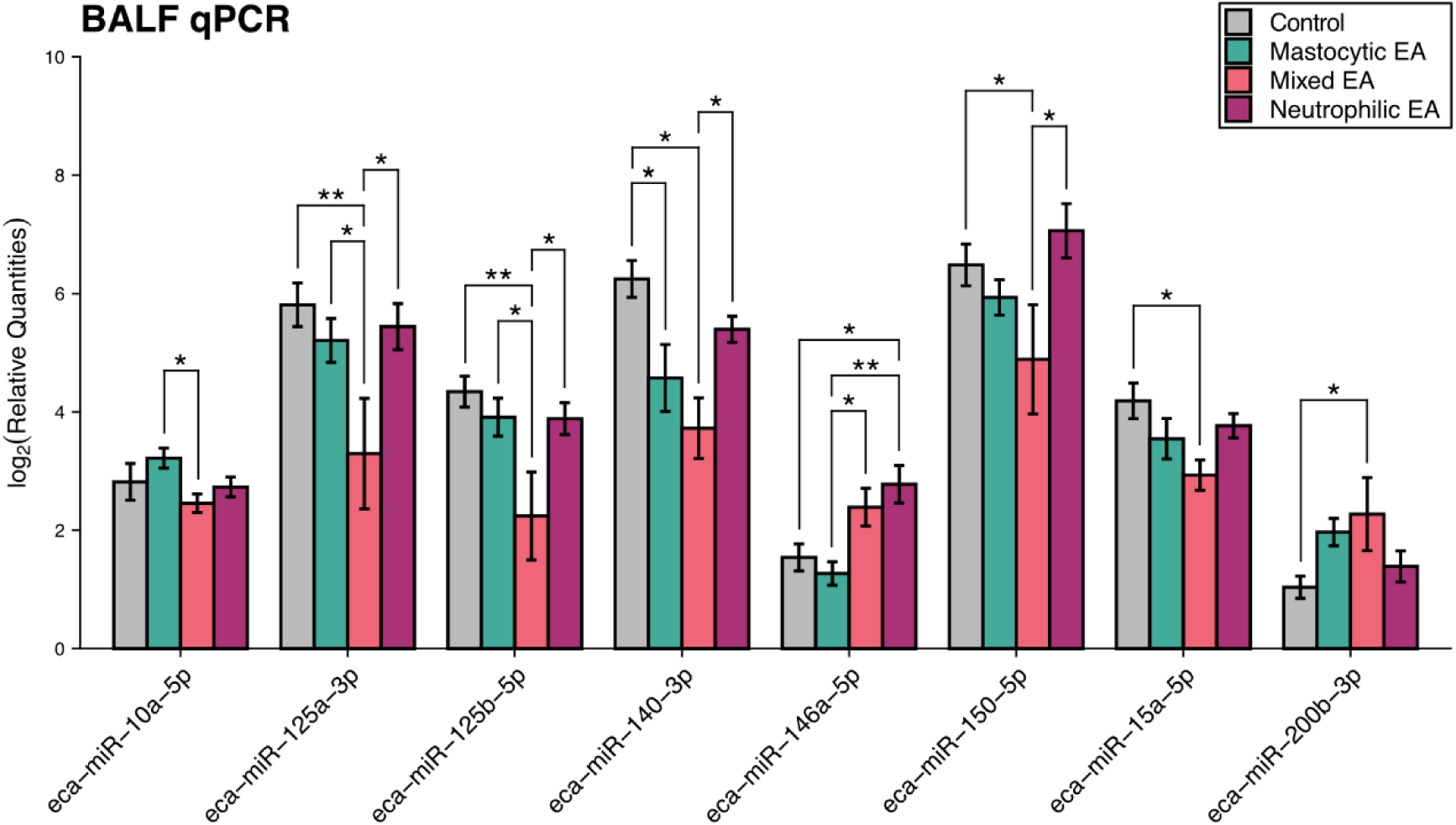
Relative quantities of significantly abundant miRNAs in BALF samples from control horses and horses with different equine asthma (EA) endotypes. Significant differences at nominal level (*P*-value < 0.05; |Fold Change| > 1.5) are marked with a star (*). Significant differences after multiple testing correction (FDR: false discovery rate < 0.05; |Fold Change| > 1.5) are marked with two stars (**).

**Table 3:** Differentially abundant miRNAs in BALF samples from control horses and horses with different equine asthma (EA) endotypes: |Fold Change| > 1.5; FDR < 0.05 (in bold) and *P*-value < 0.05; |Fold Change| > 1.5.

| Comparison | miRNA | Fold Change | *P*-value | FDR |
| --- | --- | --- | --- | --- |
| Control vs. all EA | eca-miR-140-3p | -2.708 | 0.012 | 0.37 |
|  | eca-miR-125a-3p | -2.345 | 0.03 | 0.37 |
|  | eca-miR-125b-5p | -1.976 | 0.032 | 0.37 |
| Control vs. mastocytic EA | eca-miR-140-3p | -2.578 | 0.033 | 0.809 |
| Control vs. mixed EA | **eca-miR-125a-3p** | **-6.3** | **0.001** | **0.02** |
|  | **eca-miR-125b-5p** | **-4.327** | **0.001** | **0.02** |
|  | eca-miR-140-3p | -4.768 | 0.004 | 0.052 |
|  | eca-miR-200b-3p | 2.159 | 0.031 | 0.227 |
|  | eca-miR-150-5p | -3.005 | 0.033 | 0.227 |
|  | eca-miR-15a-5p | -2.14 | 0.039 | 0.227 |
| Control vs. neutrophilic EA | eca-miR-146a-5p | 2.333 | 0.002 | 0.082 |
| Mastocytic EA vs mixed EA | eca-miR-125b-5p | -3.216 | 0.004 | 0.1 |
|  | eca-miR-125a-3p | -3.891 | 0.006 | 0.1 |
|  | eca-miR-146a-5p | 2.119 | 0.009 | 0.1 |
|  | eca-miR-10a-5p | -1.692 | 0.036 | 0.318 |
| Mastocytic EA vs. neutrophilic EA | **eca-miR-146a-5p** | **2.997** | **0.0001** | **0.002** |
| Mixed EA vs. neutrophilic EA | eca-miR-150-5p | 4.795 | 0.003 | 0.059 |
|  | eca-miR-125a-3p | 4.985 | 0.003 | 0.059 |
|  | eca-miR-125b-5p | 3.264 | 0.007 | 0.084 |
|  | eca-miR-140-3p | 2.951 | 0.043 | 0.378 |

EA: equine asthma; FDR: false discovery rate.

The miRNA eca-miR-140-3p also showed significantly reduced levels in mixed EA horses and in mastocytic EA compared to controls (**Table 3** and **Figure 3**) and was the most significantly downregulated miRNA (*P*-value=0.012) when comparing control vs. all EA endotypes combined. The other miRNA that showed high significance in the differential abundance analysis in BALF samples was eca-miR-146a-5p. This miRNA was almost three-fold more abundant in neutrophilic EA compared to the mastocytic EA group and 2.3-fold more abundant in the neutrophilic EA group compared to control horses (**Table 3**, **Figure 3** and **Supplementary Table 8**).

Moreover, similar to what was found for the eca-miR-200b-3p assay in serum (**Figure 2**), this miRNA showed elevated abundance in the mixed EA group, and also in the mastocytic EA group, but only the contrast between controls and horses with mixed EA yielded a significant result at nominal level (**Table 3** and **Figure 3**).

### Dynamics of microRNA abundance in serum and bronchoalveolar lavage fluid

To analyze the dynamics of abundance for the three miRNAs that showed the highest significant change in BALF (eca-miR-125a-3p, eca-miR-125b-5p, and eca-miR-146a-5p) we first compared their abundance profiles in BALF and serum with the patterns of immune cell counts measured in each individual horse (**Supplementary Table 1**). For eca-miR-146a-5p, a positive correlation between its relative abundance measured by qPCR in BALF and the neutrophil cell counts percentage was observed in horses diagnosed with either mastocytic or neutrophilic EA (**Figure 4** and **Supplementary Table 1**). For the mast cell counts in the BALF, such relationship was, however, inverse. Moreover, these patterns were lost when analyzing the abundance of this miRNA in serum (**Figure 5**).

**Figure 4:**
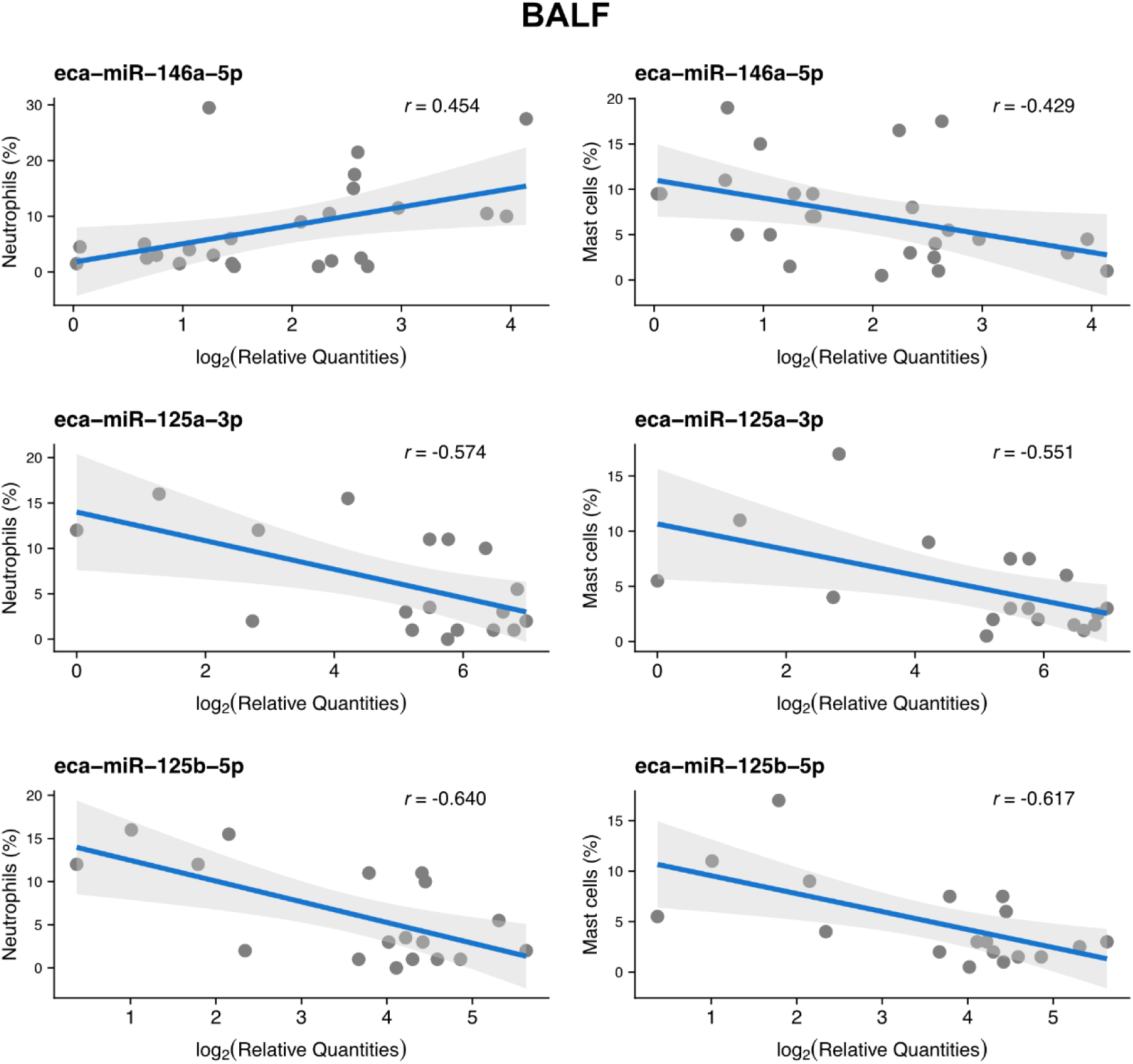
Relationship between the relative miRNA abundance of eca-miR-146a-5p, eca-miR-125a-3p, and eca-miR-125b-5p profiled by qPCR in BALF and the percentage of cells identified as neutrophils and mast cells in cytospins from horses diagnosed with either neutrophilic or mastocytic EA for eca-miR-146a-5p, and from control horses and horses diagnosed with mixed EA for eca-miR-125a-3p and eca-miR-125b-5p. A linear trend was fitted for each pairwise comparison and expressed by a Pearson’s correlation coefficient (*r*).

**Figure 5:**
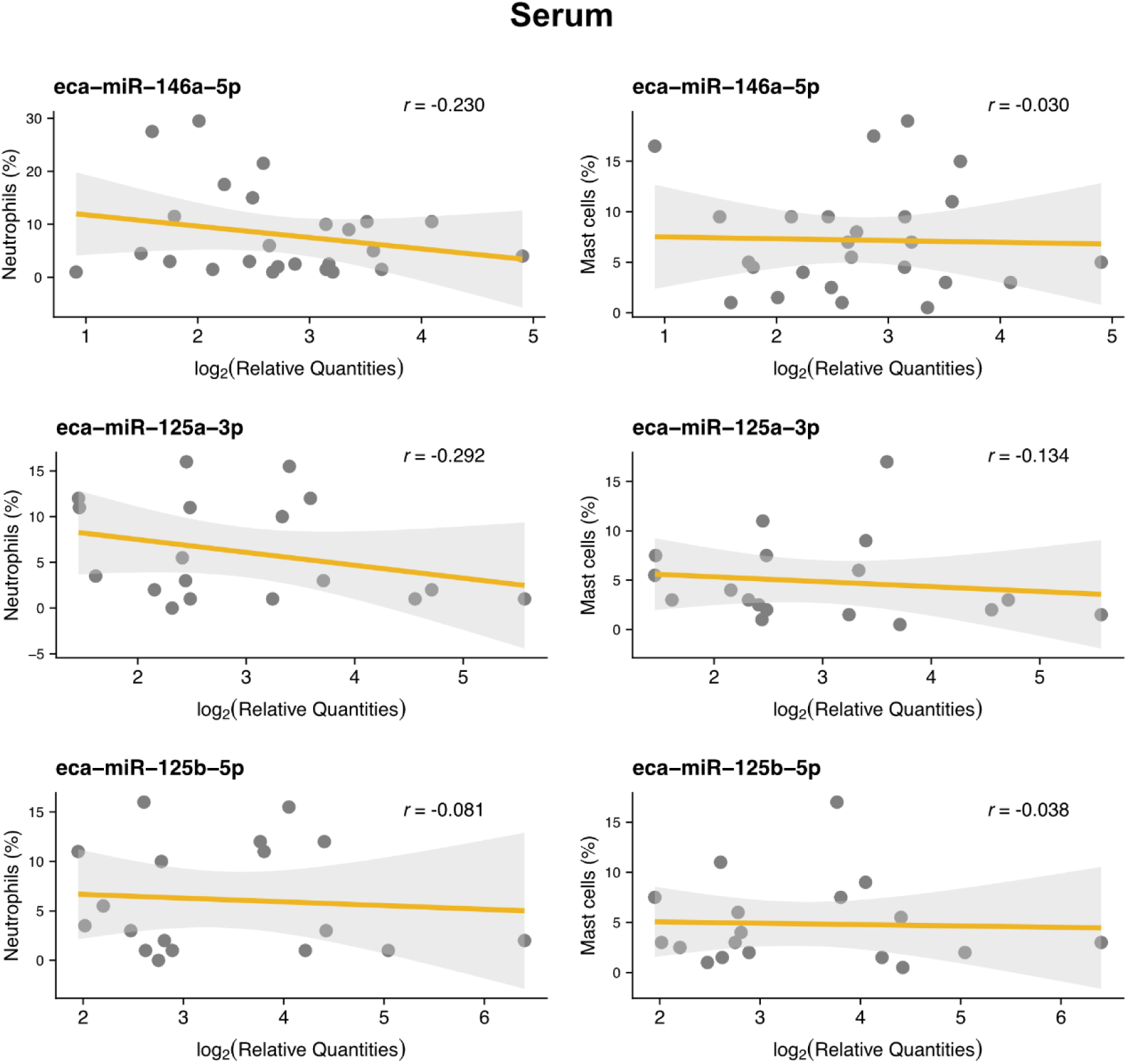
Relationship between the relative miRNA abundance of eca-miR-146a-5p, eca-miR-125a-3p, and eca-miR-125b-5p profiled by qPCR in serum and the percentage of cells identified as neutrophils and mast cells in cytospins from horses diagnosed with either neutrophilic or mastocytic EA for eca-miR-146a-5p, and from control horses and horses diagnosed with mixed EA for eca-miR-125a-3p and eca-miR-125b-5p. A linear trend was fitted for each pairwise comparison and expressed by a Pearson’s correlation coefficient (*r*).

Contrary to what was observed for eca-miR-146a-5p, both neutrophils and mast cell counts in BALF were negatively correlated with the relative abundance of eca-miR-125a-3p and eca-miR-125b-5p profiled in BALF from control horses and horses with mixed EA (**Figure 4**). Again, as seen for eca-miR-146a-5p, such a relationship was lost when using serum abundance of both miRNAs (**Figure 5**).

Besides, we compared the direction and magnitude of fold changes in serum and BALF for each EA endotype relative to controls **(Figure 6)** to assess whether miRNA abundance exhibited similar or opposing trends between the two biofluids. MiRNAs eca-miR-125a-3p and eca-miR-125b-5p had very similar curves and barely showed any change in abundance in serum reflecting the above qPCR results (**Supplementary Table 7**) but we found strongly decreased levels of these miRNAs in horses with mixed EA in BALF. For eca-miR-146a-5p, the pattern was the opposite between serum and BALF, where horses diagnosed with mixed EA and neutrophilic EA showed an increasing level of this miRNA in BALF compared to control horses, but decreased levels in serum (**Figure 6**).

**Figure 6:**
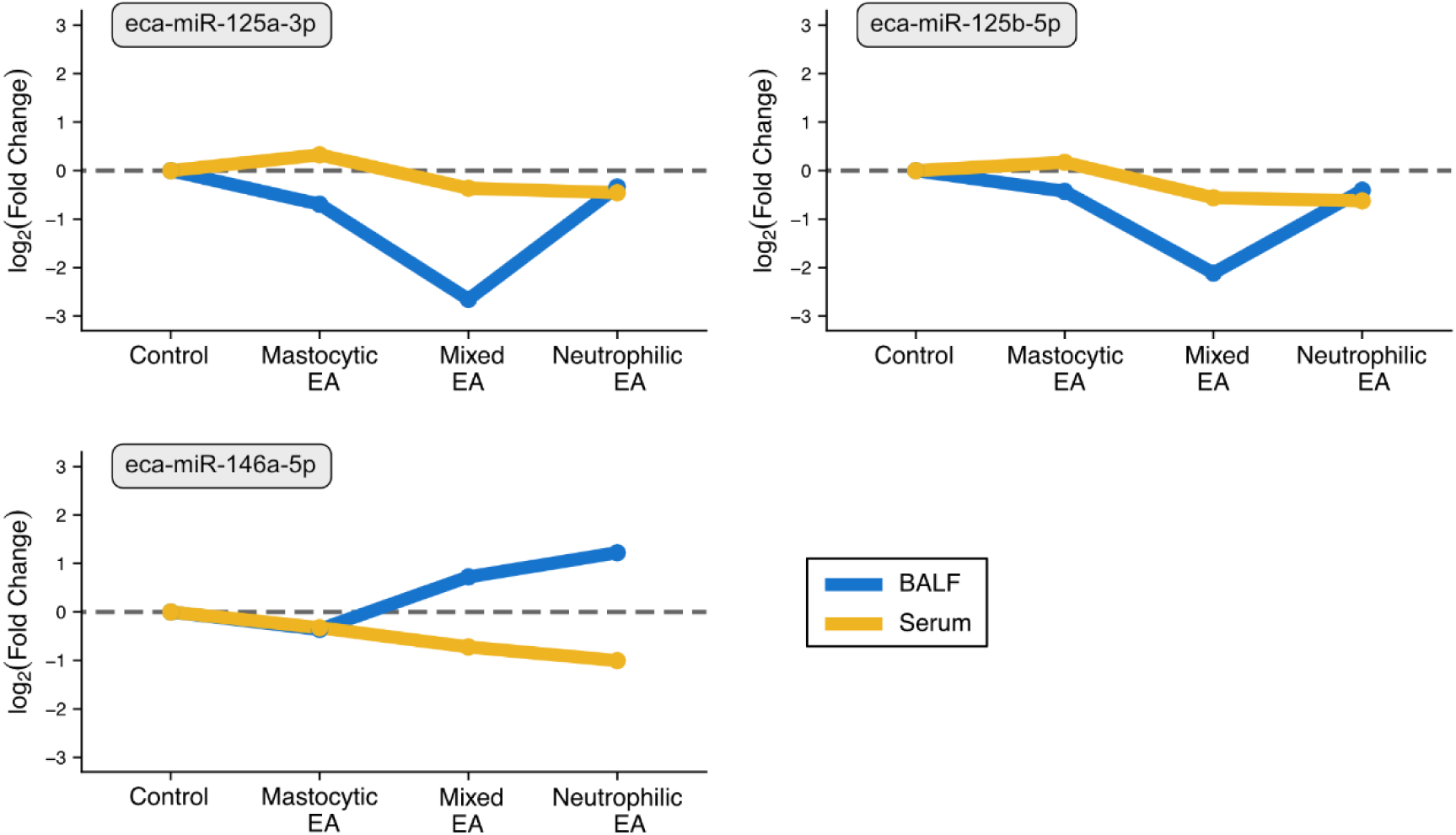
Trajectory plots across three equine asthma (EA) endotypes and control horses for the 3 miRNAs found differentially abundant in BALF. The y axis represents the log_2_FC of the qPCR results taking control horses as baseline and then comparing the profiled abundance change of the miRNA in the rest of EA groups to the control group in both serum and BALF data. If EA horses have higher abundance of the miRNA, the log_2_FC is positive, and negative if the abundance is lower.

### Target and enrichment analyses

To explore whether any of the predicted mRNA targets for the three miRNAs showing a |FC| > 1.5 and FDR < 0.05 in the differential abundance analysis (namely eca-miR-125a-3p, eca-miR-125b-5p and eca-miR-146a-5p, see **Table 3**) showed any predicted association with the pathogenesis of EA, we performed target prediction and enrichment analyses. Prediction of 8mer-targeted mRNA transcripts by any of these three miRNAs yielded a total of 2187, 3897, and 2565 binding sites for eca-miR-125a-3p, eca-miR-125b-5p, and eca-miR-146a-5p, respectively. A comprehensive list of all predicted 8mer targets is available in **Supplementary Table 9**.

Pathway enrichment analysis for all 8mer-targeted genes by any of the three miRNAs mentioned above only showed significant results for eca-miR-146a-5p (see **Table 4**).

**Table 4:** Pathway enrichment analysis (False Discovery Rate [FDR] < 0.05) from putative predicted 8mer protein-coding mRNA targets of eca-miR-146a-5p.

| Pathway | Fold Enrichment | FDR | No. Genes | Pathway Genes |
| --- | --- | --- | --- | --- |
| ErbB signaling pathway | 3.84 | 0.006 | 13 | 77 |
| Insulin signaling pathway | 3.04 | 0.006 | 17 | 127 |
| Neurotrophin signaling pathway | 2.92 | 0.027 | 14 | 109 |
| Human immunodeficiency virus 1 infection | 2.41 | 0.027 | 19 | 179 |
| MAPK signaling pathway | 2.01 | 0.047 | 24 | 271 |
| T cell receptor signaling pathway | 2.69 | 0.047 | 13 | 110 |
| Regulation of actin cytoskeleton | 2.18 | 0.047 | 20 | 208 |

Notably, several of such pathways, such as MAPK and T-cell receptor signaling, are involved in immune cell activation and airway remodeling, which are all relevant for the pathology behind EA. A full list of all enriched pathways and predicted target genes involved highlighted in our analyses is available in **Supplementary Table 10**.

## Discussion

This study is both the first to characterize BALF miRNA profiles in horses with EA and to explore differences in miRNA abundance across distinct EA cytological endotypes. We identified the miRNAs eca-miR-125a-3p, eca-miR-125b-5p, and eca-miR-146a-5p as potential EA biomarkers in BALF, demonstrating significant differential abundance between horses with EA and controls, as well as across EA endotypes.

### Comparison with previous studies of microRNAs in equine asthma

Although no previous studies have investigated miRNAs in BALF from asthmatic horses, profiling of miRNAs in lung tissue [12], airway smooth muscle [15], serum [13], and plasma neutrophil-derived exosomes [14] from horses with sEA have identified differentially abundant miRNAs, suggesting a relevant role as regulatory components in EA pathogenesis. Several of these miRNAs were also analyzed in the present study, and while some were not detected, many were profiled in BALF and/or serum. These included miR-223-3p, previously reported to be regulated in lung tissue [12]; miR-26a-5p, miR-221-3p, miR-133a-3p, miR-145-5p, and miR-21-5p, which have been shown to be regulated in airway smooth muscle [15]; miR-140-3p, previously reported to be regulated in serum [13]; and miR-155-5p and several let-7 family members, which were regulated in exosomes from plasma neutrophils [14]. However, none of these miRNAs showed significant differences between EA endotypes and controls, neither in BALF nor in serum, in the present study.

These discrepancies may reflect differences between miRNA expression within lung tissue and miRNA molecules released into the epithelial lining fluid. Additionally, previous studies focused on horses with sEA, whereas horses included in the present study predominantly had mEA. Theoretically, the miRNA response in sEA may be of greater magnitude and more readily distinguishable from controls than in mEA, which could be a contributing factor as to why we were unable to reproduce the findings from previous EA studies. Existing literature suggests that miRNA dysregulation is more pronounced locally, especially in BALF, reflecting the adjacent airway epithelial environment [29], while showing weaker concordance with systemic changes in circulating biofluids such as serum [30,31]. This is consistent with our findings, which demonstrated a stronger correlation between miRNA abundance and cellular composition in the BALF compared to our findings in serum. Furthermore, even in horses with sEA, reported effect sizes of differentially abundant miRNAs in blood have been modest, with log_2_ fold changes in serum between −0.49 and 0.47 [13], and miR-21-5p being only barely significant at nominal level in plasma neutrophil-derived exosomes [14].

### Increased abundance of eca-miR-146a-5p in BALF as a signature of neutrophilic asthma

In this study, a significant nearly three-fold increase of eca-miR-146a-5p was observed in BALF from horses with neutrophilic EA compared to the mastocytic endotype. In addition, eca-miR-146a-5p showed a positive correlation with neutrophil cell count in BALF and a negative correlation with the mast cells count. Collectively, this suggests that the regulation of this miRNA may be linked to neutrophil-driven airway inflammation. Indeed, miR-146a-5p is a key regulator of innate immunity and the NF-κB pathway in other species [32]. In humans, hsa-miR-146a exerts anti-inflammatory effects by controlling immune cell proliferation and inhibiting excessive inflammatory responses [32–34]. Interestingly, reduced hsa-miR-146a-5p has been linked to higher neutrophil-attracting chemokines in human neutrophilic asthma [35]. The increased level of the miRNA in the BALF found in neutrophilic EA in this study may thus reflect a compensatory anti-inflammatory response, with eca-miR-146a-5p being upregulated in epithelial lining fluid to counteract excessive inflammation, a mechanism also reported in humans [36].

Underscoring these findings, target prediction and pathway analysis for eca-miR-146a-5p indicated regulation of several inflammation-related pathways: i) MAPK signaling, which controls proliferation, differentiation, apoptosis, and stress responses [37,38]. MAPKs synergize with NF-κB, promoting neutrophilic inflammation, and the p38 MAPK endotype is specifically implicated in airway inflammation in asthma [39,40]. ii) T-cell receptor (TCR) signaling, which drives T-cell proliferation, differentiation, and cytokine production, while also engaging NF-κB, contributing to neutrophilic responses in both EA and human asthma [41,42]. EA involves Th1, Th2, and Th17 responses [8,12], reflecting TCR-mediated naïve T-cell differentiation.

Eca-miR-146a-5p was also of interest in the study by Vargas et al. [14], where miRNAs in plasma neutrophil-derived exosomes from EA were evaluated. In that study eca-miR-146a-5p was not differentially abundant when comparing the exosomes’ miRNA content between sEA and control horses, which is in line with what we found in serum samples for this miRNA. Thus, while in our study miR-146a-5p appears to be increased locally in association with neutrophilic inflammation, systemic levels in blood may not mirror this pattern. However, it should be noted that in both studies a (non-significant) decrease in serum abundance of this miRNA was observed in EA horses compared to controls.

### Reduced levels of eca-miR-125a-3p and eca-miR-125b-5p in BALF as a signature of mixed asthma

In horses with mixed EA profiles, characterized by elevated mast cells and neutrophils, eca-miR-125a-3p and eca-miR-125b-5p levels were significantly lower compared to controls (fold changes −6.3 and −4.33). Further, a negative correlation was found between both the neutrophil and mast cell count in the BALF and the abundance of both miRNAs. This suggests they may serve as a molecular signature of mixed inflammatory EA. To the authors’ knowledge, no EA studies have examined these miRNAs. The miR-125b-5p has recognized anti-inflammatory roles. In humans, it is downregulated in bronchial biopsies from asthma patients, correlating with severe airflow obstruction and elevated neutrophils [43], and low levels are reported in smokers [44] and asthmatic children [45]. Intranasal mmu-miR-125b in murine asthma reduced IL-4 and IL-13, goblet cell differentiation, and mucus production [45]. Mechanistically, it targets the 3′UTR of TNF-α, reducing protein levels and inflammation [32,46]. While miR-125a-3p’s immune functions are less defined, it can act as a tumor suppressor [47] and modulates host defense via targeting the 3’UTR of the UV radiation resistance-associated gene (*UVRAG)* [32]. Besides, its downregulation may enable effective immune responses [48].

Overall, the reduced eca-miR-125a-3p and eca-miR-125b-5p levels likely reflect a dysregulated inflammatory environment in the mixed EA endotype, though further studies are needed to clarify their extracellular roles in horses.

### Considerations on sampling serum and BALF in asthma

When comparing the two sample types used in this study, BALF and serum each offer distinct advantages and limitations as matrices for miRNA biomarker discovery. BALF provides direct access to the cellular and extracellular environment of the lower airways, better reflecting local lung pathology, including asthma-related changes in miRNA expression [49]. BALF-derived miRNAs (particularly those contained in cells or extracellular vesicles) have been shown in human asthma to discriminate patients from controls and to associate with airway inflammatory and remodeling pathways, supporting their relevance as lung-specific biomarkers [50]. In contrast, serum captures the circulating miRNA pool and reflects systemic physiology, integrating signals from multiple organs, including but not limited to the lungs [51,52]. In the present study, the strongest disease-associated miRNA signals were detected in BALF, where the miRNAs eca-miR-125a-3p, eca-miR-125b-5p, and eca-miR-146a-5p emerged as promising candidates for novel BALF EA biomarkers. Notably, human studies have implicated hsa-miR-125b-5p and hsa-miR-146a-5p in asthma-related airway inflammation and clinical severity, further supporting their biological relevance as biomarkers in the equine model [13,43].

### Study limitations

The EA endotype assignment of the horses analyzed in the present study relies solely on BALF cytology, for which variability has been observed and described in previous literature [7]. Moreover, the method used for profiling miRNAs in this study was qPCR, which is a hypothesis-driven technique that restricts detection to a predefined *ad hoc*-selected list of candidate miRNAs. Therefore, other potentially relevant or novel miRNAs involved in EA cannot be identified using this method. In contrast, hypothesis-free approaches such as small-RNA sequencing would enable comprehensive profiling of all miRNAs present, allowing detection of both known and novel miRNAs to assess differences in their abundance across EA endotypes.

## Conclusions

This study identified eca-miR-125a-3p, eca-miR-125b-5p, and eca-miR-146a-5p as promising biomarkers in BALF for distinguishing EA endotypes. Eca-miR-125a-3p and eca-miR-125b-5p were significantly downregulated in BALF from horses with mixed EA, whereas eca-miR-146a-5p showed differential abundance, being downregulated in serum but upregulated in BALF of horses with neutrophilic EA. Pathway analysis indicated that eca-miR-146a-5p regulates immune-relevant pathways, including MAPK and T-cell receptor signaling, highlighting its role in airway inflammation. These findings support the potential of miRNAs as molecular biomarkers to complement or enhance cytology-based diagnostics for EA and strengthen our understanding of the EA endotypes and pathogenesis. Validation in independent cohorts and integration with sequencing approaches are needed to confirm these candidates and uncover additional relevant miRNAs before attempting clinical application.

## Acknowledgements

The authors would like to thank Minna Jakobsen, Tina B. N. Mahler and Ilona E. Baranski for excellent technical support in the qPCR part of this study. We would also like to thank the technical staff at the Large Animal Teaching Hospital, University of Copenhagen, for their assistance with sample collection and preparation, especially Tina Roust and Maria Rhod for their help in the laboratory. The Project was funded by “The Grant for Interdisciplinary Master Thesis projects”, School of Veterinary Medicine & Animal Sciences, University of Copenhagen to SC and SH. EM-S acknowledges the financial support from the Villum Fonden (Villum Experiment project no. 57875).

